# Stepping into a dangerous quagmire: environmental determinants of human-lancehead pit vipers (*Bothrops* genus) contact resulting in injuries, Brazilian Amazon

**DOI:** 10.1101/329649

**Authors:** João Arthur Alcântara, Paulo Sérgio Bernarde, Jacqueline Sachett, Ageane Mota da Silva, Samara Freire Valente, Henry Maia Peixoto, Marcus Lacerda, Maria Regina Oliveira, Ivan Saraiva, Vanderson de Souza Sampaio, Wuelton Marcelo Monteiro

## Abstract

Despite significant and successful efforts in Brazil regarding snakebites in the areas of research, antivenom manufacture and quality control, training of health professionals in the diagnosis and clinical management of bites, little is known about determinants of snakebites incidence in order to further plan interventions to reduce the impact of this medical condition. Understanding the complexity of ecological interactions in a geographical region is important for prediction, prevention and control measures of snakebites. The aim of this investigation is to describe spatial distribution and identify environmental determinants of human- lancehead pit vipers (*Bothrops* genus) contact resulting in injuries, in the Brazilian Amazon. Aggregated data by municipality was used to analyze the spatial distribution of *Bothrops* bites cases and its relationship with geographic and environmental factors. Eight geo-environmental factors were included in the analysis as independent variables: (1) tree canopy loss increase; (2) area with vegetation cover; (3) area covered by water bodies; (4) altitude; (5) precipitation; (6) air relative humidity; (7) soil moisture; and (8) air temperature. Human- lancehead pit vipers (*Bothrops* genus) contact resulting in envenomings in the Amazon region is more incident in lowlands [-0.0006827 (IC95%: −0.0007705; - 0.0005949), p<0.0001], with high preserved original vegetation cover [0.0065439 (IC95%: 0.0070757; 0.0060121), p<0.0001], with heaviest rainfall [0.0000976 (IC95%: 0.0000925; 0.0001026), p<0.0001] and higher air relative humidity [- 0.0081773 (IC95%: −0.0107681; −0.0055865), p<0.0001]. This association is interpreted as the result of the higher forest productivity and abundance of pit vipers in such landscapes.

**Author summary:** Despite successful efforts in Brazil regarding snakebites in the areas of research, antivenom manufacture and quality control and training of health professionals, little is known about determinants of snakebites incidence in order to further plan interventions to reduce the impact of this medical condition. Understanding the complexity of ecological interactions in a geographical region is important for prediction, prevention and control measures of snakebites. The aim of this study is to describe spatial distribution and identify environmental determinants of human- lancehead pit vipers (*Bothrops* genus) contact resulting in injuries, in the Brazilian Amazon. An increase in the forest productivity with a higher availability of some types of prey, such as frogs and amphibians, anurans and lizards, was suggested as a cause for the higher snake abundance in the rainy season. Probably due to the higher forest productivity and abundance of pit vipers in such landscapes, human-lancehead pit vipers contact resulting in envenomings in the Amazon region is more incident in lowlands, with high preserved original vegetation cover, with heaviest rainfall and higher air relative humidity.

## Background

The neotropical pitviper clade of *Bothrops* and *Bothrocophias* is distributed throughout South America and associated continental islands, and includes species that range into Central America, Mexico, and the Caribbean (1–4) Commonly known as lanceheads, the group comprises 47 species allocated in discrete species groups (4–9). Within *Bothrops*, several species groups have been repeatedly recovered and named: *Bothrops alternatus* group, *Bothrops neuwiedi* group, *Bothrops jararaca* group, *Bothrops atrox* group and *Bothrops taeniatus* group (9). Species of *Bothrops* occupy all main ecosystems, from rainforests to grasslands and other dry habitats (4,10). This genus includes mostly terrestrial species (e.g. *B. alternatus* and *B. neuwiedi*), as well as many that use vegetation, from the semi-arboreal *B. jararaca* (11) to the almost completely arboreal *B. bilineatus* (12–14). In at least some species, such as *B. atrox*, juveniles are found frequently on the vegetation than adults (15–17). In the Brazilian Amazon and surrouding *cerrado* areas, there are 12 species of pit vipers, belonging to the *Bothrops* and *Bothrocophias* genera. Five of them (*Bothrops lutzi*, *B. marmoratus*, *B. mattogrossensis*, *B. moojeni* and *B. pauloensis*) are present only in *cerrado* areas, while the others are characteristic of the Amazon rainforest environments (*Bothrops atrox*, *B. bilineatus*, *B. brazili*, *B. marajoensis*, *B. taeniatus*, *Bothrocophias hyoprora* e *B. microphthalmus*) (4,18). Major species responsible for *Bothrops* envenomings in the Brazilian Amazon and surrouding cerrado areas are psented in Figure 1.

**Figure 1.**
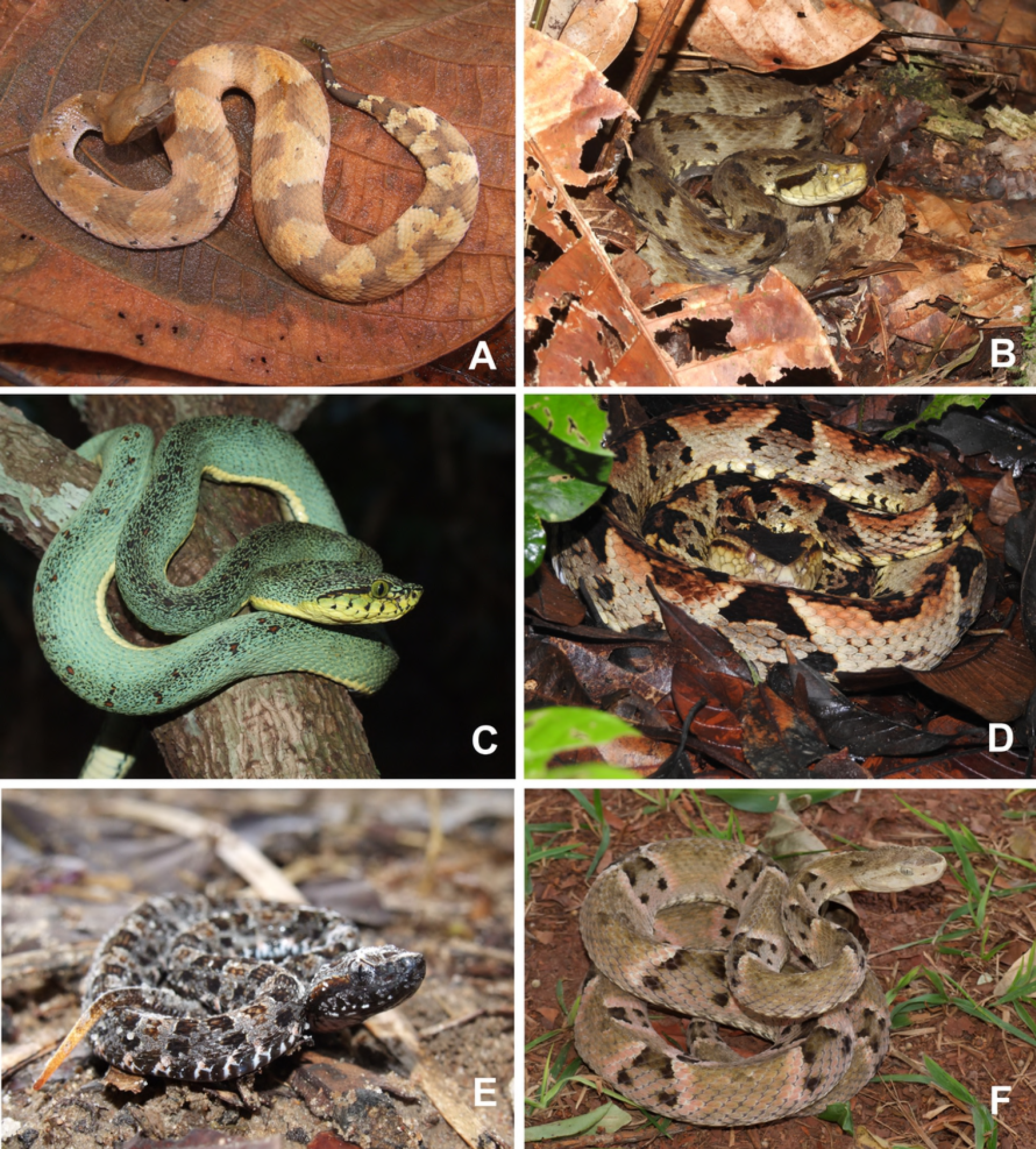
Pictures of some species responsible for *Bothrops* envenomings in the Brazilian Amazon and surrouding cerrado areas. a) *Bothrocophias hyoprora*; b) *Bothrops atrox*; c) *Bothrops bilineatus bilineatus*; d)*Bothrops brazili*; e) *Bothrops moojeni*; f) *Bothrops mattogrossensis*.”

*Bothrops atrox*, the Amazonian lancehead, inhabits mostly forests, although it may be occasionally found in disturbed habitats around human settlements, including deforested areas (pastures and crops) and in urban environments (4,16,18–21) This snake is the responsible for more cases and fatalities in the Amazon than any other venomous snakes, causing 80-90% of the snake envenomings in the region (22). This species has predominantly nocturnal activity; the adult hunts preferentially on the ground, while the young are mostly found on the vegetation (16,17), presenting a generalist diet, preying on centipedes, fish, anurans, lizards, other snakes, birds and small mammals (16,23–28). Despite the wide geographic distribution in the Amazon, *B. atrox* venoms share the same family of toxins, as PIII and PI snake venom metalloproteinase, phospholipase A2, serine proteinase, cysteine-rich secretory protein, L-amino acid oxidase and C-type lectin-like (29,30). A PI metalloproteinase, called batroxase, isolated from *B. atrox* venom, has fibrinolytic, thrombolytic activities and induces weak bleeding through the digestion of the extracellular matrix components such as laminin, type IV collagen and fibronectin (31). *Bothrops* envenoming shows pain, swelling, regional lymphadenopathy, ecchymosis, blistering and necrosis as the most common local clinical manifestations (32–34). Secondary bacterial infections were observed in around 40% of the *Bothrops* snakebites (34). Spontaneous systemic bleeding and acute renal failure are common systemic complications after *Bothrops* envenomings (32–34).

In the Brazilian Amazon, snakebites are under the influence of precipitation, likely because snakes in the Amazon exhibit increased activity during months with heaviest rainfall. Moreover, in this period, the snakes look for upland areas during flooding, which increases the likelihood of contact between humans and snakes (35,36). The contact between snakes and human populations is often associated with extractive or agricultural activities, which can increase snakebite burden in conditions of land management or deforestation. Urban growth provides changes in the natural habitat of snakes, which can lead to a greater probability of snakebites. Despite significant and successful efforts in Brazil regarding snakebites in the areas of research, antivenom manufacture and quality control, training of health professionals in the diagnosis and clinical management of bites, little is known about determinants of snakebites incidence in order to further plan interventions to reduce the impact of this medical condition. Understanding the complexity of ecological interactions in a geographical region is important for prediction, prevention and control measures of snakebites. The aim of this investigation is to describe spatial distribution and identify environmental determinants of human-lancehead pit vipers (*Bothrops* genus) contact resulting in injuries, in the Brazilian Amazon.

## Methods

### Ethical clearance

This study was approved by the Ethics Review Board (ERB) of the *Núcleo de Medicina Tropical* of the *University of Brasîlia* (approval number 1.652.440/2016).

### Study design, data source and definitions

An ecological study design was carried out including the states of Acre, Amapá, Amazonas, Mato Grosso, Pará, Rondônia, Roraima, Tocantins and Maranhão, whose ecotypes are classified in the Amazon biome. The study area occupies 5,016,136.3 km^2^, corresponding to about 59% of the Brazilian territory and has a population of more than 24 million people (37). The entire 775 second administrative level subdivisions, called municipalities, were defined as units of analysis. The cartography also used municipalities as unit of analysis, performed with QGIS (Version 2.18.17 LTR). The dependent variable for mapping was the snakebite incidence, presented as the mean absolute number of cases per year, reported from 2010 to 2015, using municipality population as the denominator, standardizing per 100,000 inhabitants. Data on the municipal populations was obtained from the 2010 official census and the intercensus projections (38). *Bothrops* envenomings in humans are officially reported to the Brazilian Ministry of Health. The department responsible for snakebites surveillance provided the data presented here (Supplementary File 1). Although *Bothrops* contact with humans resulting in envenomings generally is diagnosed based on the clinico-epidemiological profile, the positive predictive value of this type of identification reaches 97.8-100% compared to immunoassay techniques using monoclonal antibodies, due to the high prevalence of this species perpetrating injuries (32,34); such a high value can be interpreted as indicating an excellent accuracy and internal validity of the dependent variable used in this study. Although only the number of cases has been used for the analyses, all database variables were checked for duplicates and completeness by two independent researchers before analysis. The general characteristics of the cases, such as gender, age, anatomical region of the injury, area of occurrence, work-related injury, schooling, ethnical background, time elapsed from the bite until medical assistance and outcome were described. The rainfall climatology was used to construct seasonality maps (39).

Aggregated data by municipality was used to analyze the spatial distribution of *Bothrops* bites cases and its relationship with geographic and environmental factors. Eight geo-environmental factors were included in the analysis as independent variables: (1) tree canopy loss increase; (2) area with vegetation cover; (3) area covered by water bodies; (4) altitude; (5) precipitation; (6) air relative humidity; (7) soil moisture; and (8) air temperature.

### Definitions

The variables in this study were defined as follows:

#### Tree canopy loss

Average annual deforested area in the municipalities between 2007 and 2014, which was measured by the average annual percentage of the municipal area that lost forest vegetation; estimated based on the computer assisted analysis of a series of images from Lansat, Cbers, UK-2-DMC or Resourcenet. The analysis is performed by TerraLib/TerrAmazon project. The detection of deforested area, vegetation cover and cloud area are used to estimate the total increment of deforested area as described in PRODES methodology, considering the automatically detected area plus the estimated area under cloud cover, according the *Coordenadoria Geral de Observação da Terra Programa Amazônia* (PRODES) (40);

#### Area with vegetation cover

Percent (%) of municipal area covered by vegetation in 2010, automatically detected by the image processing as described in PRODES methodology (40);

#### Area covered by water bodies

Percent (%) of municipal area covered by water bodies in 2010, automatically detected by the image processing as described in PRODES methodology (40);

#### Altitude

Measured as the lowest point within a county in meters above mean sea level using the global digital elevation model geo-processed by *Agência Nacional de Aviação Civil* (41);

#### Precipitation

Defined as the deposition of water to surface of Earth, in the form of rain, snow, ice or hail. Is measured in millimeter (mm) and one millimeter of rain corresponds to 1 liter per square meter of water on the surface. In this study, we used the accumulated value in the period evaluated. This data were compose from Unified Precipitation Project that are underway at NOAA Climate Prediction Center (CPC), every day with spatial resolution of 0.5° latitude × 0.5° longitude at surface level (42);

#### Air relative humidity

Defined as the ratio, expressed in percent, of the amount of water vapor in a given volume of air to the amount that this volume could contain if the air were saturated. (43); This data were compose from National Centers for Environmental National Centers for Environmental Prediction (NCEP) reanalysis, every 6 hours (0 to 18) with spatial resolution of 2.5° latitude × 2.5° longitude at surface level (43);

#### Soil moisture

Defined as the water that is maintained in the spaces between soil particles (cm^3^ water/cm^3^ soil), in other words the water is available in the upper layer of soil. This data were compose from National Centers for Environmental National Centers for Environmental Prediction (NCEP) reanalysis, every 6 hours (0 to 18) with spatial resolution of 2.5° latitude × 2.5° longitude between 0 - 10 cm in soil (43);

#### Temperature

Defined as a quantity of heat that exists in the air and measured in degree Celsius (°C). This data were compose from National Centers for Environmental National Centers for Environmental Prediction (NCEP) reanalysis, every 6 hours (0 to 18) with spatial resolution of 2.5° latitude × 2.5° longitude at 2 meters of the surface (43).

### Data analysis

The variables were entered individually into a univariate logistic regression model and preselected if p≤0.20. Subsequently, variance inflation factor (VIF) was estimated to verify the relationship between all preselected independent variables (check for potential collinearity), in which coefficient >10 were considered high. For this study none VIFs were higher than 10. Interactions between biologically plausible variables were examined (rain vs. temperature; area with vegetation cover vs. canopy tree loss and air relative humidity vs. precipitation), if found significant (p<0.05), interaction terms were kept for further analysis. The eight geo-environmental factors were included in the analysis as independent continuous variables. A Poisson model was also used to estimate the regression coefficient between snakebites incidence and the geo-environmental factors. Multivariable models were built in a backward elimination step, resulting in a final model in which only variables with p<0.05 were kept. The goodness-of-fit of the final model was tested using Hosmer-Lemeshow, p>0.05. In addition, an adjusted model was constructed by including three variables as potential confounders into the model, with equal weights: (1) Human Population Density by municipality, assuming that snakebites are density-dependent, i.e., the contact rate between human and *Bothrops* individuals depends upon the local population density (44); and (2) Access to Health System and (3) Health System Effectiveness, subcomponents of the Mean Health System Performance Index (MHSPI) (45). Statistical analyses were performed using the STATA statistical package version 13 (Stata Corp. 2013).

## Results

### Exploratory and descriptive analysis of the snakebites

According to the official reporting system, 70,816 snakebites were recorded in the Amazon Region in the study period. From this total, 13,442 cases (19.0%) were not included in the analysis because their classification as non-venomous bites (4,886 cases), *Lachesis* bites (5,217 cases), *Crotalus* bites (3,103 cases) or *Micrurus* bites (236 cases).

The total of cases eligible in this study was 57,374 *Bothrops* snakebites, resulting in an incidence rate of 37.2 cases per 100,000 person/year. There was a slight variation in the annual incidence rates during the study period. Incidence was higher in 2011 (38.8 per 100,000 person/year), and lower in 2015 (36.2 per 100,000 person/year). All the variables retrieved from the original dataset presented completeness higher than 70% (Table 1). Most of the snakebites occurred in males (45,091 cases; 78.6%). Regarding the area of occurrence, 86.6% were reported in rural areas. The most affected age group was between 18 and 45 years old (31,568 cases; 55.0%). Admixed population was the most reported in the ethnicity field (40,499 cases; 74.4%). The most affected education group was the group with ≤4 years of schooling (18,276 cases; 44.1%). A proportion of 40.1% of the snakebites were related to work activities. Most of the snakebites occurred in the lower limbs (83.9%). Regarding time elapsed from the bite until medical assistance, 78.3% of the cases received treatment within the first six hours after the snakebite, 16.4% within 6–24 hours and 5.3% with more than 24 hours after bite.

**Table 1.**
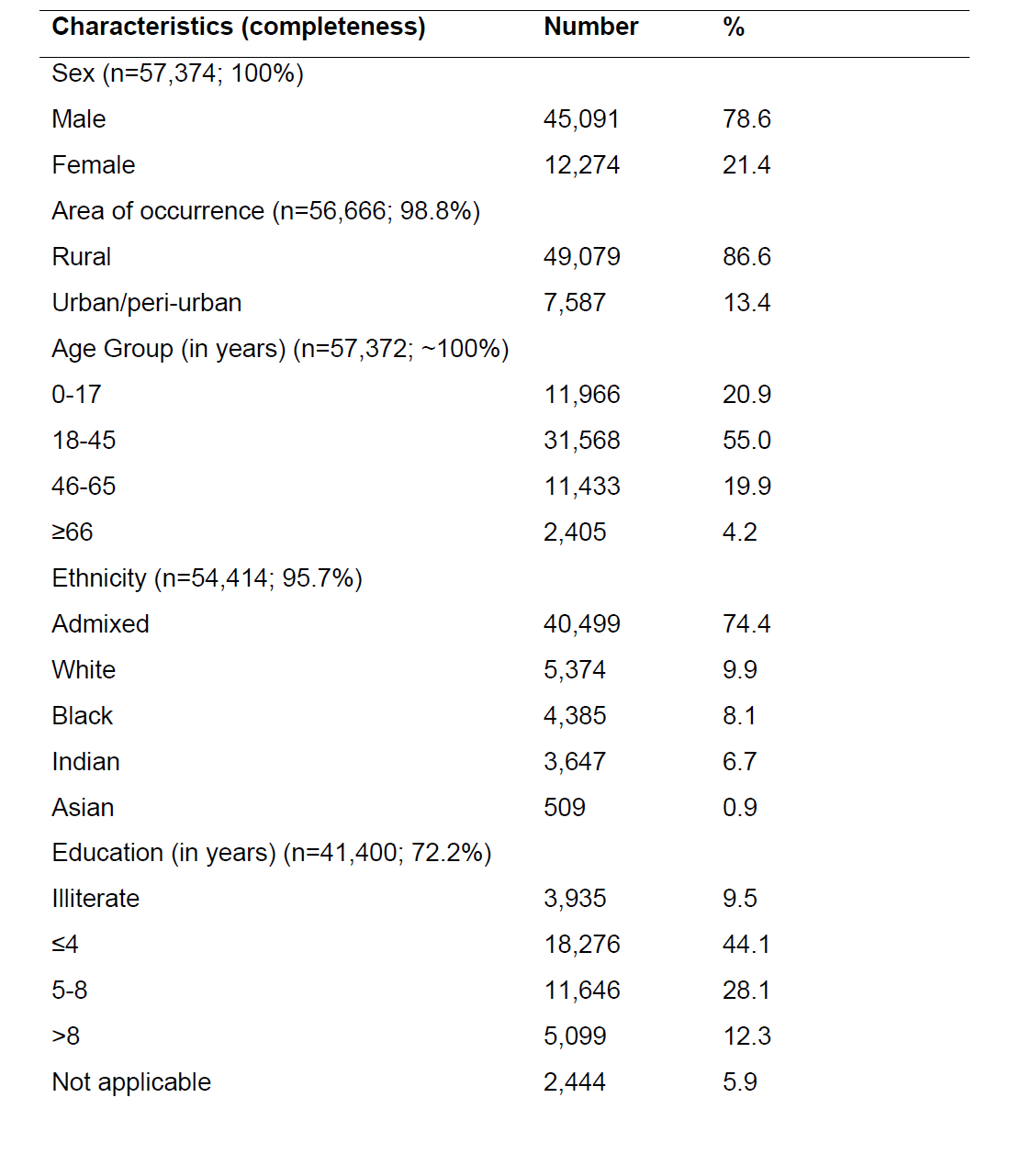
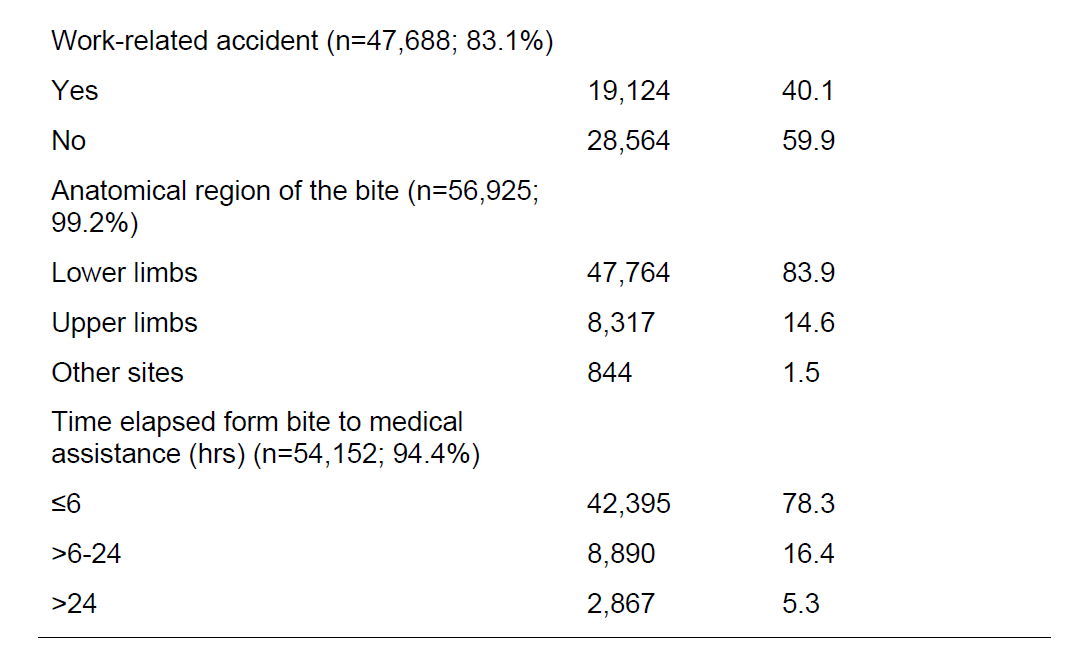
Epidemiological characteristics of patients who are victims of *Bothrops* snakebites, Brazilian Amazon, from 2010 to 2015.

| Characteristics (completeness) | Number | % |
| --- | --- | --- |
| Sex (n=57,374; 100%) |  |  |
| Male | 45,091 | 78.6 |
| Female | 12,274 | 21.4 |
| Area of occurrence (n=56,666; 98.8%) |  |  |
| Rural | 49,079 | 86.6 |
| Urban/peri-urban | 7,587 | 13.4 |
| Age Group (in years) (n=57,372; ~100%) |  |  |
| 0-17 | 11,966 | 20.9 |
| 18-45 | 31,568 | 55.0 |
| 46-65 | 11,433 | 19.9 |
| ≥66 | 2,405 | 4.2 |
| Ethnicity (n=54,414; 95.7%) |  |  |
| Admixed | 40,499 | 74.4 |
| White | 5,374 | 9.9 |
| Black | 4,385 | 8.1 |
| Indian | 3,647 | 6.7 |
| Asian | 509 | 0.9 |
| Education (in years) (n=41,400; 72.2%) |  |  |
| Illiterate | 3,935 | 9.5 |
| ≤4 | 18,276 | 44.1 |
| 5-8 | 11,646 | 28.1 |
| >8 | 5,099 | 12.3 |
| Not applicable | 2,444 | 5.9 |
| Work-related accident (n=47,688; 83.1%) |  |  |
| Yes | 19,124 | 40.1 |
| No | 28,564 | 59.9 |
| Anatomical region of the bite (n=56,925; 99.2%) |  |  |
| Lower limbs | 47,764 | 83.9 |
| Upper limbs | 8,317 | 14.6 |
| Other sites | 844 | 1.5 |
| Time elapsed form bite to medical assistance (hrs) (n=54,152; 94.4%) |  |  |
| ≤6 | 42,395 | 78.3 |
| >6-24 | 8,890 | 16.4 |
| >24 | 2,867 | 5.3 |

The nine states reported cases during the 6-year study period. The highest incidence rates were reported in the states of Pará (57.0 per 100,000 person/year), Roraima (53.3 per 100,000 person/year) and Tocantins (49.7 per 100,000 person/year). The lowest incidence rates were reported in the states of Maranhão (15.9 per 100,000 person/year), Rondônia (22.6 per 100,000 person/year) and Amazonas (29.1 per 100,000 person/year). Intermediate incidence rates were reported in the states of Mato Grosso (32.0 per 100,000 person/year), Amapá (34.9 per 100,000 person/year) and Acre (41.1 per 100,000 person/year). Several municipalities had incidence rates higher than 100 cases per 100,000 peron/year, distributed unevenly in the study area. The Northwest of the State of Amazonas, North of Roraima, North of Pará on the border with Amapá and central part of the state of Tocantins, show spots of higher incidence rates (Figure 2). The municipality of Alto Alegre, in the State of Roraima, presented the highest rate among all municipalities (358.3 per 100,000 person/year), followed by the municipalities of Anajás (338.9 per 100,000 person/year) and Afuá (303.7 per 100,000 person/year), in the State of Pará.

**Figure 2.**
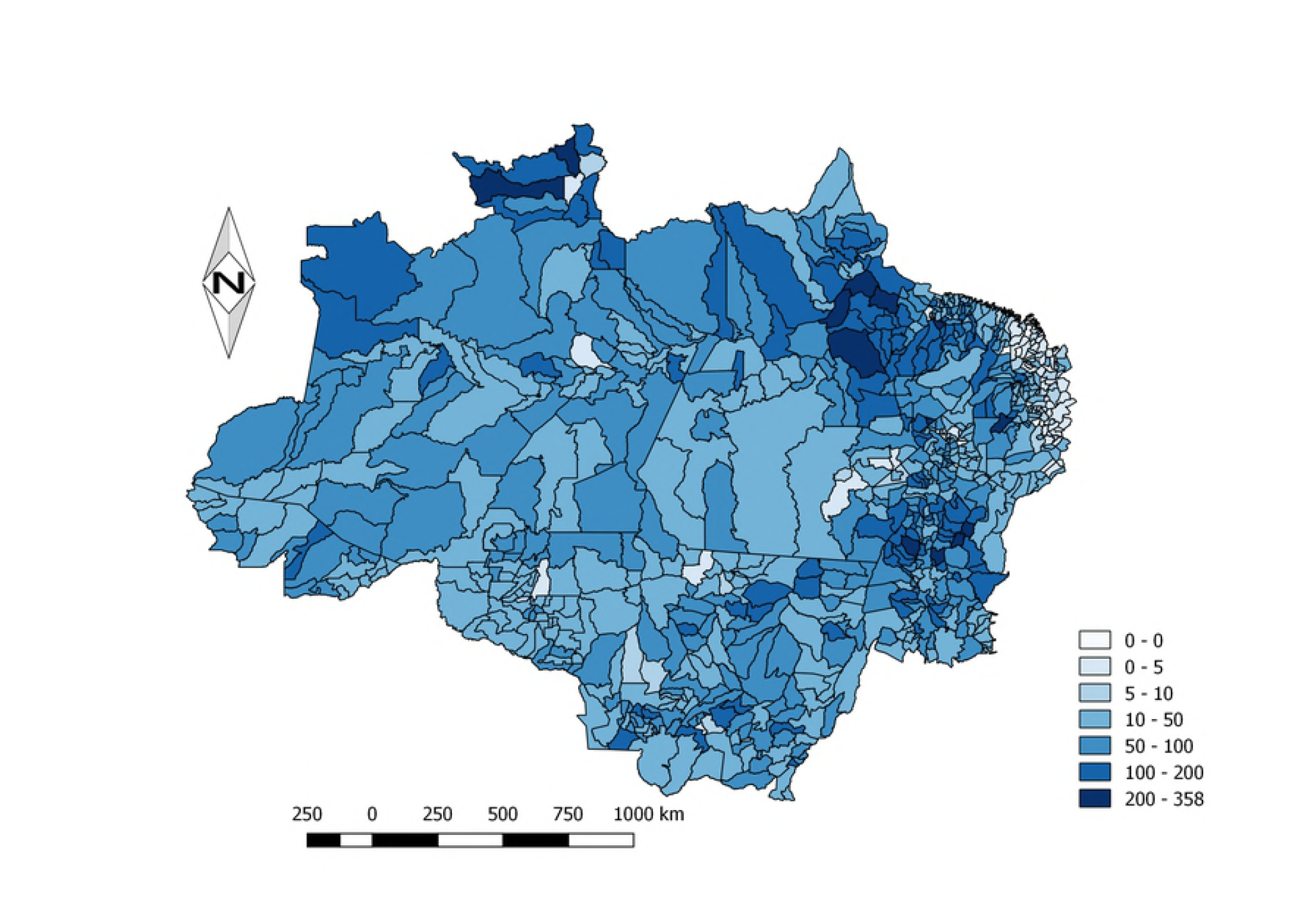
Spatial distribution of *Bothrops* snakebites in the Brazilian Amazon from 2010 to 2015. Map were created using incidence per 100,000 inhabitants. Snakebites are largely distributed in the Amazonian states, with several counties presenting incidences higher than 100 cases per 100,000 inhabitants. The Northwest of the State of Amazonas, North of Roraima, North of Pará on the border with Amapá and central part of the state of Tocantins, show spots of higher incidence rates.

### Descriptive analysis of geo-environmental variables

Mean tree canopy loss in the study area was 35.6% (±33.4%), being lower in Western Brazilian Amazon, namely in the North and West of the state of Amazonas, West of the state of Acre and Amapá. A higher tree canopy loss is observed in the state of Maranhão, in the extreme East of the states of Pará and Acre, and in some municipalities of the state of Mato Grosso and Rondônia. The mean area with vegetation cover is 25.31% (±27.4%), being lower in the states of Tocantins, Maranhão and in the South of the state of Mato Grosso. The mean area covered by water bodies is 2.9% (±6.9%) and is larger in the municipalities located in the banks of the Solimões, Negro and Amazonas rivers and in the Pantanal region of the state of Mato Grosso. Regarding altitude, the whole study area had a mean of 153.7 meters above sea level (±148.6 meters). Altitude is lower in municipalities located in the banks of the Amazonas river and higher in the states of Mato Grosso, Tocantins, south of the Maranhão, peaking in the north of the state state of Roraima. The average value of accumulated in the study area is 9,356.2 millimeters (±2,335.1 millimeter), ranging from <5,000 millimeters in some municipalities in the periphery of the region to >13,000 millimeters in the extreme Western Amazon. Mean air relative humidity in the study area is 82.9% (±6.7%), ranging from ˜70% in municipalities Southern Amazon border to >95% in the Western Amazon. Mean soil moisture in the study area is 0.3270047 cm^3^ water/cm^3^ soil (±0.0280811 cm^3^/cm^3^), being lower in the states of Maranhão, Tocantins and Mato Grosso and higher in the state of Amazonas. Mean air temperature in the region is 25.2°C (±0.81°C), ranging from ˜23°C in the state of Roraima to >27°C in the extreme Eastern Amazon (Figure 3).

**Figure 3.**
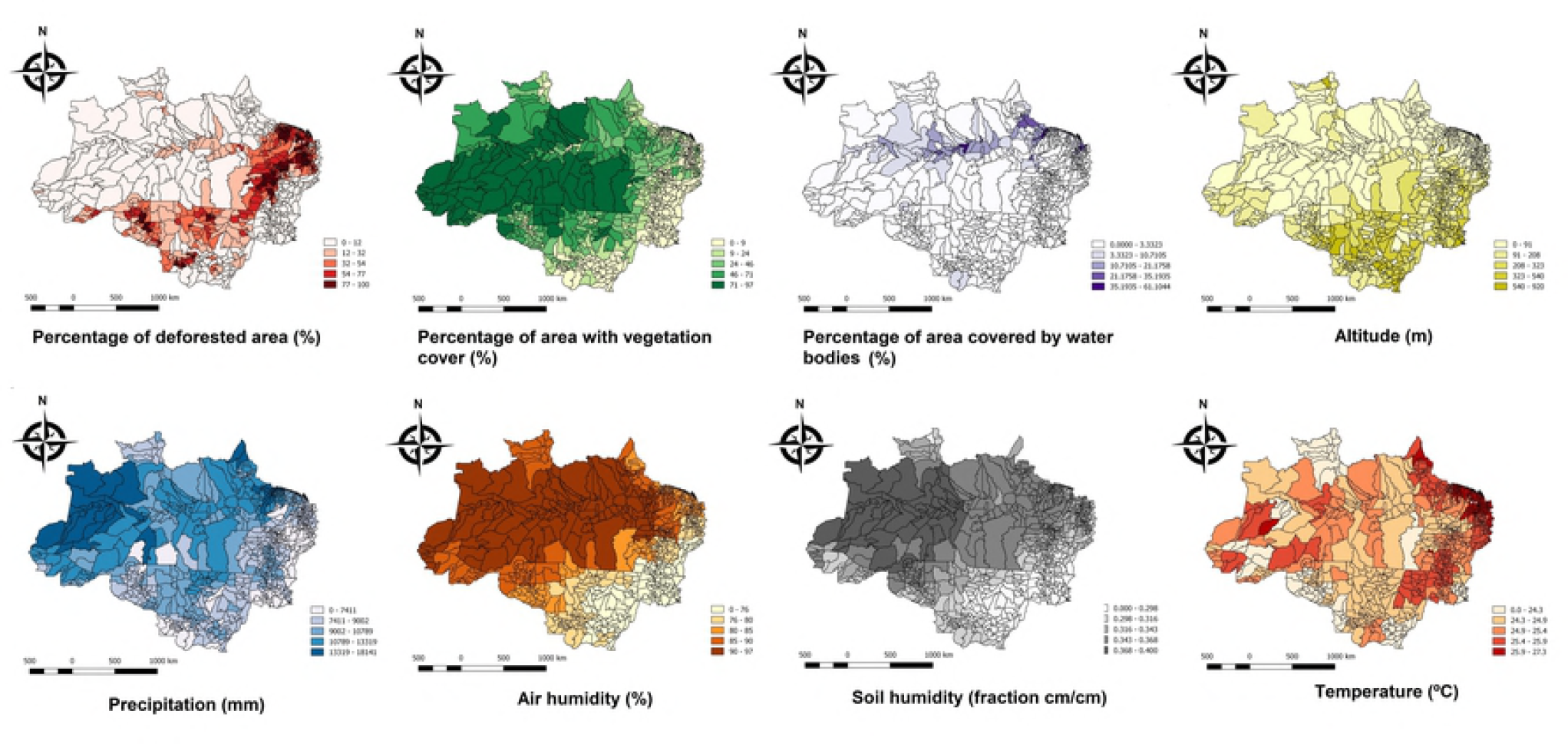
Descriptive analysis of geo-environmental variables. A) Tree canopy loss increase. B) Area with vegetation cover. C) Area covered by water bodies. D) Altitude. E) Precipitation. F) Air relative humidity. G) Soil Moisture. H) Air temperature.

### Univariate analysis and correlation test

Tree canopy loss increase [-0.0042463 (IC95%: −0.0062533;-0.0022393), p<0.0001] and air temperature [-0.1101975 (IC95%: −0.1919551;-0.0284399), p=0.008] were variables negatively related to pit vipers contact rate. Percentage of vegetation cover [0.0022327 (IC95%: 0.0003861; 0.0040793), p=0.018] and precipitation [0.0000721 (IC95%: 0.0000483; 0.000096), p<0.0001] were variables positively related to pit vipers snakebite incidence. Percentage of water bodies cover [-0.0003366 (IC95%: (−0.011507;0.0108337), p=0.953], altitude [-0.0003323 (IC95%: −0.0008188; 0.0001542), p=0181], air relative humidity [0.0084803 (IC95%: −0.0007394; 0.0176999), p=0.071] and soil moisture [0.0036591 (IC95%: (−0.0175959; 0.024914), p=0.736] were not significantly related to snakebite incidence rates in the univariate analysis (Table 2).

**Table 2.**
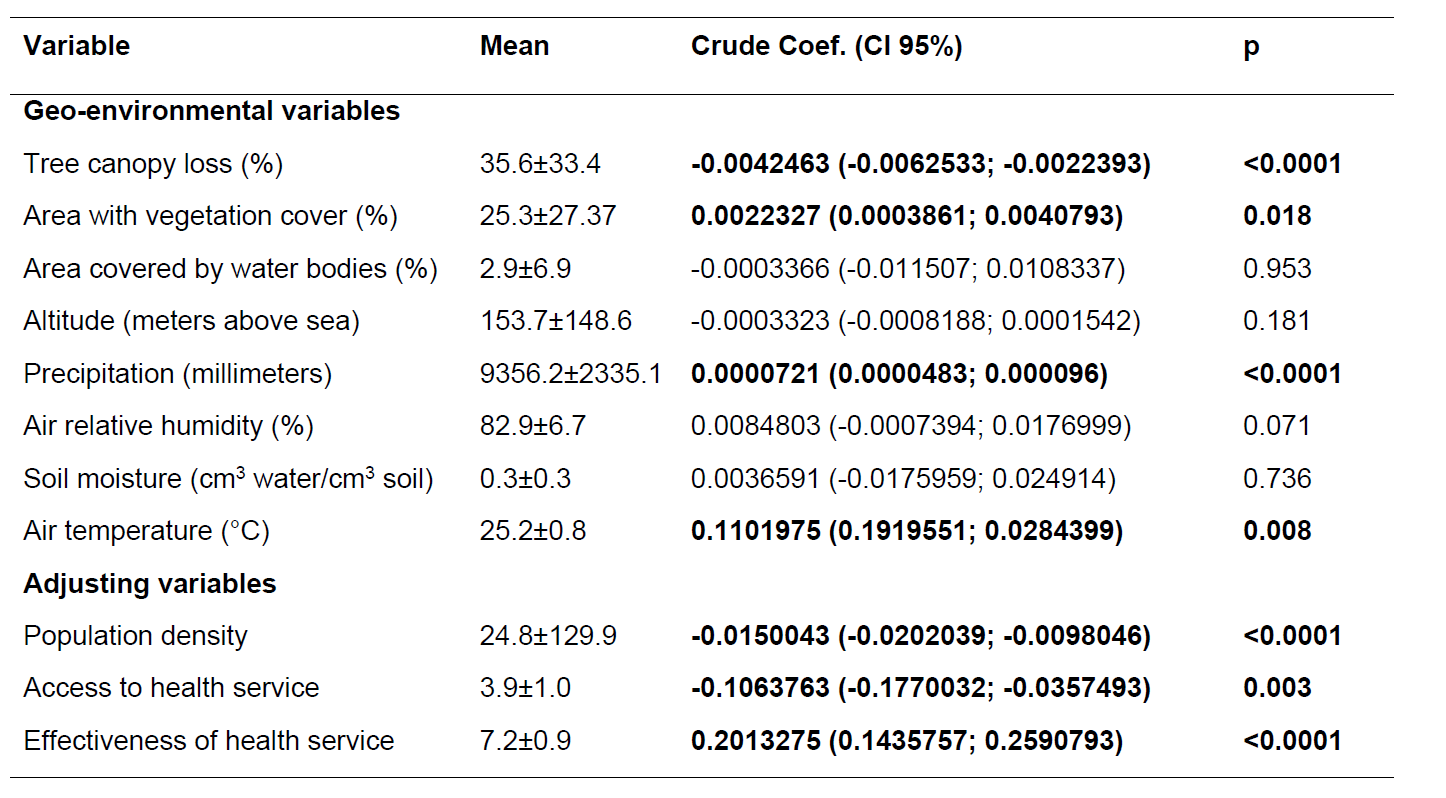
Association of geo-environmental variables with *Bothrops* snakebite incidence rates in the Brazilian Amazon in the univariate analysis.

| Variable | Mean | Crude Coef. (CI 95%) | p |
| --- | --- | --- | --- |
| **Geo-environmental variables** |  |  |  |
| Tree canopy loss (%) | 35.6±33.4 | **-0.0042463 (-0.0062533; -0.0022393)** | **<0.0001** |
| Area with vegetation cover (%) | 25.3±27.37 | **0.0022327 (0.0003861; 0.0040793)** | **0.018** |
| Area covered by water bodies (%) | 2.9±6.9 | -0.0003366 (-0.011507; 0.0108337) | 0.953 |
| Altitude (meters above sea) | 153.7±148.6 | -0.0003323 (-0.0008188; 0.0001542) | 0.181 |
| Precipitation (millimeters) | 9356.2±2335.1 | **0.0000721 (0.0000483; 0.000096)** | **<0.0001** |
| Air relative humidity (%) | 82.9±6.7 | 0.0084803 (-0.0007394; 0.0176999) | 0.071 |
| Soil moisture (cm<sup>3</sup> water/cm<sup>3</sup> soil) | 0.3±0.3 | 0.0036591 (-0.0175959; 0.024914) | 0.736 |
| Air temperature (°C) | 25.2±0.8 | **0.1101975 (0.1919551; 0.0284399)** | **0.008** |
| **Adjusting variables** |  |  |  |
| Population density | 24.8±129.9 | **-0.0150043 (-0.0202039; -0.0098046)** | **<0.0001** |
| Access to health service | 3.9±1.0 | **-0.1063763 (-0.1770032; -0.0357493)** | **0.003** |
| Effectiveness of health service | 7.2±0.9 | **0.2013275 (0.1435757; 0.2590793)** | **<0.0001** |

Human population density, access to health system and health system effectiveness were tested in an univariate model to assess their role as potential confounders. Population density [-0.0150043 (IC95%: −0.0202039;-0.0098046), p<0.0001] and access to health service [-0.1063763 (IC95%; −0.1770032;- 0.0357493), p=0.003] were negatively related to snakebites incidence rate. Health system effectiveness [0.2013275 (IC95%: 0.1435757;0.2590793), p<0.0001].

There was a notable seasonality of *Bothrops* bites over the year, with a pronounced increase of cases in the rainiest trimester across the study area (Figure 4).

**Figure 4.**
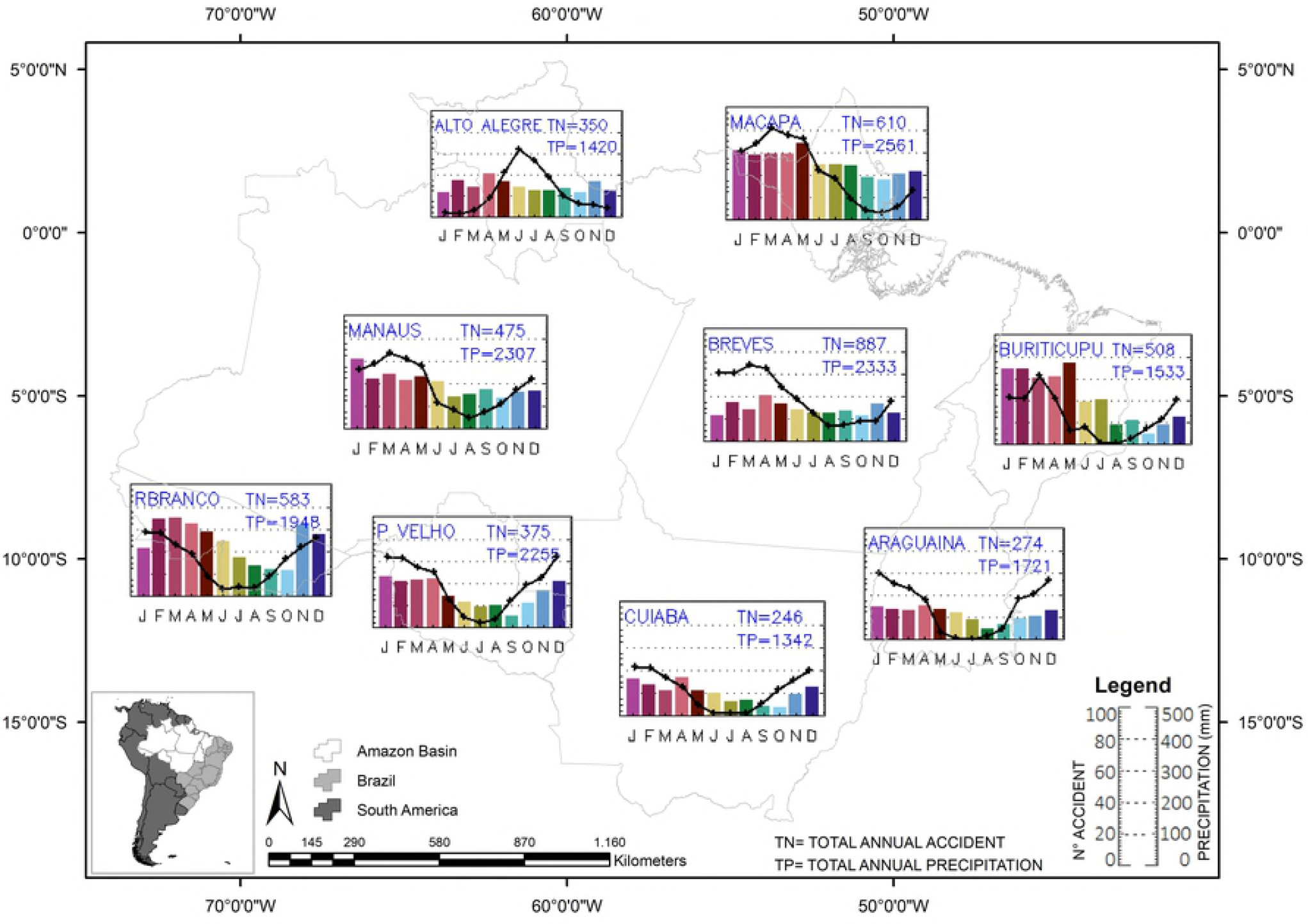
Univariate analysis and correlation test. Chart of correlation graph between the geo-environmental variables with *Bothrops* bites incidence, as well as their respective values of p and R. A) Tree canopy loss increase. B) Area with vegetation cover. C) Area covered by water bodies. D) Altitude. E) Precipitation. F) Air relative humidity. G) Soil Mosture. H) Air temperature.

### Multivariable analysis

After adjustment by human population density, access to health system and health system effectiveness, altitude [-0.0006827 (IC95%: −0.0007705; −0.0005949), p<0.0001] and air temperature [-0.160476 (IC95%: −0.1759117;-0.1450403), p<0.0001] were negatively related to pit vipers contact rate. Moreover, percentage of vegetation cover [0.0065439 (IC95%: 0.0070757; 0.0060121), p<0.0001], precipitation [0.0000976 (IC95%: 0.0000925; 0.0001026), p<0.0001] and air relative humidity [-0.0081773 (IC95%: −0.0107681; −0.0055865), p<0.0001] were variables positively related to *Bothrops* bite incidence (Table 3).

**Table 3.** Association of geo-environmental variables with *Bothrops* snakebite incidence rates in the Brazilian Amazon in the multivariate analysis.

| Variable | Adjusted Coef. [Final model] (CI 95%) | p |
| --- | --- | --- |
| Geo-environmental variables |  |  |
| Tree canopy loss (%) | **0.0001072 (-0.0002437; 0.0004582)** | **0.549** |
| Area with vegetation cover (%) | **0.0065439 (0.0070757; 0.0060121)** | **<0.0001** |
| Altitude (meters above sea) | **-0.0006827 (-0.0007705; -0.0005949)** | **<0.0001** |
| Precipitation (millimeters) | **0.0000976 (0.0000925; 0.0001026)** | **<0.0001** |
| Air relative humidity (%) | **0.0081773 (0.0107681; 0.0055865)** | **<0.0001** |
| Air temperature (°C) | **-0.160476 (-0.1759117; -0.1450403)** | **<0.0001** |

**Legend:** Adjusted by human population density, access to health system and health system effectiveness.

## Discussion

In the Brazilian Amazon, although urban population predominates, most of the snakebites were recorded in adult males living in rural areas. This epidemiological profile was reported previously in the Amazon (35,36), and probably is related to a significant higher exposition of this population group when exerting agricultural or forestry activities, in the habitat mostly selected by lancehead pit vipers. Several studies have reported on habitat use by *B. atrox*, the easiest snake to find in comparison to other species in Central Amazonia (46), and concluded that the species is mainly found on the ground or climb into understory vegetation, in the case of juveniles (15–17). In this study, most of the snakebites occurred in the lower limbs, suggesting that adult snakes than juveniles, are the major responsible by the human injuries, inflicted after the contact of human individuals, barefoot or with lower limbs not fully protected, with snakes using the ground to rest (especially during the day), hunt and move (15,46). Some agroforestry activities developed in the Amazon are very conducive to contact of workers with snakes, such as the Brazil nut and palm tree fruits harvest, in which scouring the leaf litter for the collection of fallen fruits enables the frequent finding of *B. atrox*, *B. taeniatus* and *B. brazili* (47,48).

Information on habitat use and activity in *B. atrox* is helpful to understand of the biology of this widespread and ecologically important species in the Amazon. Unfortunately, abundance and use of habitats of pit vipers in different ranges of anthropization are not properly studied. In this work, percentage of vegetation cover was positively related to *Bothrops* bite incidence, indicating that the original rainforest environment maintenance is crucial for population density in levels that warrant an intense contact rate between *Bothrops* individuals and humans. In this preserved environment, snakes probably find more propitious terrestrial (leaf litter) and occasionally arboreal microhabitats where these animals use the space for survival, especially in relation to prey availability (15–17,49). Populations from a variety of taxonomic groups, including snakes and their preys are declining in response to the loss of early-successional habitats (50). Although very less frequent, reports of urban cases are noteworthy, showing that pit vipers are able to occupy a wide environmental gradients and maintain populations in forest fragments within very anthropized areas, as observed for *B. atrox* in the Amazon (18), *B. moojeni* and *B. neuwiedi* in Cuiabá, Southern Amazon (51) and *B. jararaca* in Southern Brazil (52). Furthermore, Amazonian cities outskirts are often encrusted in sylvatic environments, enabling snake populations to frequent the peridomiciliary areas where rodents, marsupials, frogs and other synanthropic animals serve as food sources (51). Forestry, hunting and recreational activities are common in this urban relictual forested areas or deeper in the forests along urban fringes are expect to be risk factors for snakebites (49).

In this investigation, significant higher snakebite incidence rates were found in areas with higher precipitation indexes. As previously flagged in other studies, most of the snakebites in the Amazon are recorded in the rainy season, with some geographical variations in the region (35,49,53,54). Some authors suggest that this seasonal pattern is related to the flooding in areas of land adjacent to streams margins, the location of the riverine villages, forcing the snakes to upland areas, increasing the likelihood of contact between humans and snakes (35). From another perspective, it is well established that snakes in the Amazon exhibit increased activity during months with higher rainfall, including *Bothrops atrox* and *B. bilineatus* (13,14,16,17). In the region of Manaus, Central Amazon, *B. atrox* present an unimodal seasonal activity in summer reflected in more productive collections of specimens in this season (16). Cunha and Nascimento (19) also reported on seasonal activity in *B. atrox* in eastern Amazonia (decrease from June-November). Thus, the results may apply for other Amazonian localities as well, and explain the notable seasonality of *Bothrops* bites over the year, with a pronounced increase of cases in the rainiest trimester across all the study area, as shown in this study. Furthermore, in the Amazon the incidence of juveniles also occurred mainly during the rainy months (13,16). Recruitment in this period can be related to the higher availability of prey (16) and/or due to other environmental factors (e.g., increase in humidity and temperature) (17,55). An increase in the forest productivity with a higher availability of some types of prey, such as frogs and amphibians, anurans and lizards, was suggested as a cause for the higher snake abundance in the rainy season (16,17,56,57). Interestingly, a positive relationship between amount of rainfall and the number of snake stomachs containing anurans was found in tropical snakes (56). Rainfall, which is highly correlated with air humidity is often suggested as an important factor determining the seasonal incidence of tropical snakes (16,17,56,58,59), consistent with the positive relation between air relative humidity and *Bothrops* bite incidence found here.

Within the environmental variables, rainfall is the most discriminative for the tropical region, since other functions such as air temperature and relative humidity are are contidioned by the first (60,61). In the Amazon basin, climate is indicated as a function of the rainy and dry season and an average alternative is made around 2300 mm.year-1, reaching a number of 3,500 mm.year-1 in the northwest of the region (62). In this context, relative air humidity is directly modulated by rainfall and presents higher values during the rainy season of each region, but as the largest varieties occur in the northern Legal Amazon (state of Roraima), with minimum values around of the 30% (60,62). In this study, locations with the highest air humidity had a high incidence of accidents. On the other hand, air temperature was negatively affected by the snakebite rates, that is, the higher the temperature, the lower incidence of snakebites. This probaly occurs because the air is heated by the incident energy on the terrestrial surface, in long wave form (sun radiation), which in turn backs up energy in long wave form and heats the air (60,61). In the Amazon region, the incident radiation flux is constant throughout the year, which is able to influence small changes in temperature over the year, except for the forest region that causes cooling incidence during winter in the Southern Hemisphere (63). Incidence was higher in 2011 and lower in 2015, coinciding with the El Niño-Southern Oscillation, tied to variation in the amount and distribution of precipitation and river water levels in the Amazon basin (64).

Herein, municipalities located in the extensive lowlands of the Amazon basin presented high snakebite incidence rates. These plain area is surrounded by plateaus, located between the Guianas plateau, the Brazilian plateau and the Atlantic ocean, in which incidence was significantly lower. Although the seasonally flooded lowland (called ‘ *várzeas*’) occupies only a small part of this region, extending along the banks of the Amazon River and its tributaries, the vast expanses of low-plateaus or low-sedimentary plateaus (called ‘mainland‘) are saved from common floods. However, sedimentary plateaus are the origin of numerous streams flowing into the ‘*várzeas*’ area watercourses and lakes. In this landscape, in summary, water dividers present a slightly higher altitude in relation to the parafluvial catena. These lowlands enclosed within a forest canopy are the most likely ecosystem for encountering snakes of the *Bothrops* genus (15,48). Snake richness on follows four distinct patterns: decreasing, low-elevation plateaus, low-elevation plateaus with mid-elevation peaks, and mid-elevation peaks. In general, it is expected that elevational reptile richness and snakebites are most strongly correlated with temperature, mediated by precipitation, and decreases on all high landscapes (48;65-67). Consistently, in the Central Amazon, the distribution of *B. atrox* is not uniform within the forest, with a density of about 6.4 times higher near streams, probably due to the increase in prey availability (15). As indigenous and riverside populations live in general in human settlements in the river banks, this riverscape encompasses ecological processes that conduce humans to contact snakes.

Record keeping may have been influenced by the nature of the surveillance system. Some patients with mild bites in inaccessible areas may not be reported to hospitals and those evolving to severity may die on the way before reaching medical attention. To minimize this possible bias, environmental correlations were adjusted by *Access to Health System and Health System Effectiveness*, subcomponents of the *Mean Health System Performance Index* (MHSPI) (45). These variables were included to adjust the possible differences in sensitivity of the surveillance case definition among municipalities across the study area, i.e., the ability of the epidemiological surveillance to identify all possible *Bothrops* snakebites in the community, depending mostly on the access of the individuals bitten to the health services and the effectiveness of this system do diagnose and report the cases properly (68).

In conclusion, human-lancehead pit vipers (*Bothrops* genus) contact resulting in envenomings in the Amazon region is more incident in lowlands, with high preserved original vegetation cover, with heaviest rainfall and higher air relative humidity. This association is interpreted as the result of the higher forest productivity and abundance of pit vipers in such landscapes.

## Acknowledgments

We would like to thank the Brazilian Ministry of Health, for providing database used in this study.

## Supporting Information Legends

**S1 Table**. Study database.

